# A Triple-Measure Model for Estimating Cranial–Brain Size Across Age Groups

**DOI:** 10.64898/2026.10.12.758588

**Authors:** Cicero Moraes, Johari Yap Abdullah, Angela Beatriz Pomatti, Andrej Shbat, Jiří Šindelář, Ivo Klepáček, Elena Varotto, Lis Moura, Thiago Beaini, Francesco Maria Galassi

## Abstract

**Background:** Head circumference is widely used as an indirect indicator of brain growth and cranial capacity; however, its isolated use may lead to imprecise interpretations, particularly in borderline cases of microcephaly and megalencephaly. Complementary screening approaches that remain accessible and non-invasive may therefore be useful, especially in settings with limited access to neuroimaging.

**Objective:** To develop and evaluate an experimental, non-invasive model for estimating endocranial volume based on a combined cranial metric, termed the Triple-Measure, and to compare its performance with head circumference alone across different age ranges.

**Methods:** One hundred forensic facial approximations with known cranial scale were analyzed. Head circumference, sagittal arc, and coronal arc were measured on three-dimensional skull models, and their sum was defined as the Triple-Measure. Endocranial volume was obtained through direct segmentation of computed tomography data or by anatomical deformation of donor models. Regression analyses and validation procedures were performed, and the results were contextualized using literature-derived datasets to assess compatibility with established population parameters.

**Results:** The Triple-Measure showed a stronger correlation with endocranial volume (R = 0.83; R² = 0.69) than head circumference alone (R = 0.76; R² = 0.57), a difference confirmed by an F-test (F ≈ 17.03, p < 0.001). Validation analyses performed on independent CT cases demonstrated good agreement and consistency between predicted and measured values. The sample distribution was compatible with large population datasets derived from the literature. Based on these findings, an analog lookup table and a digital calculation tool were developed to support exploratory application across adult and developmental stages, with improved sensitivity in borderline situations.

**Conclusions:** The Triple-Measure may provide a more informative anthropometric proxy for endocranial volume than head circumference alone while remaining non-invasive and accessible. Given its exploratory nature, the method is best interpreted as a complementary screening approach intended to support clinical decision-making and to indicate when further imaging evaluation may be warranted. Additional studies with larger clinical datasets are recommended to refine its applicability across populations.

**Highlights:**

- Triple-measure shows improved association with endocranial volume compared to head circumference
- Proposed as a non-invasive, low-cost screening approach for micro- and megalencephaly
- Analog lookup table and digital calculator developed for exploratory and practical use
- Framework extended across neonatal, pediatric, and adult age groups using integrated datasets

## Introduction

### Anthropometry and Craniometry

Anthropometry is the science dedicated to the measurement of human body proportions. The term derives from the Greek *ánthr*ō*pos* (man) and *métron* (measure), and its initial conception was more linked to the idealization of the body than to the systematic quantification of its variations [1]. Although present in classical art and architecture, it was only from the 17th century, with the medical studies of Elsholtz, that anthropometry began to acquire scientific contours. Its consolidation, however, occurred in the 19th century with Lambert Adolphe Quetelet, who introduced the application of statistical methods to the study of the human body, using Gauss’s normal curve and developing the concept of the “average man” [2].

In the 20th century, anthropometry evolved with the standardization of techniques, such as those from the International Society for the Advancement of Kinanthropometry (ISAK), in which head circumference measurement follows a precise protocol: an inextensible tape measure is used, positioned over the glabella (the most prominent point between the eyebrows) and the opisthocranion (the most posterior part of the skull), with the child in the Frankfurt position to ensure alignment. The measurement, recorded in centimeters, is repeated two or three times for greater reliability [3].

This technique is essential for monitoring brain growth, which reaches about 84% of adult size at 12 months, reflecting the rapid synaptic development and myelination in the first year of life. At 2 years, head circumference reaches 90% of adult size, with slower growth after 5 years [3]. Beyond growth tracking, the measurement serves as a reliable indicator of neurological and nutritional development: deviations may signal microcephaly, associated with cognitive deficits or infections such as Zika, or macrocephaly, linked to hydrocephalus or genetic syndromes. Regular monitoring combined with weight and height measurements is therefore crucial for early interventions, enabling timely referral for medical evaluation when deviations are identified [3].

The head circumference is used to construct growth curves, comparing measurements with World Health Organization (WHO) percentiles. Children with values between the 3rd and 97th percentiles are considered normal, while deviations (below the 3rd or above the 97th) require medical evaluation. In the school environment, these curves allow for the early identification of developmental problems, such as cognitive delays or nutritional deficits, which can impact adulthood. For example, early malnutrition, which slows cranial growth, is associated with lower cognitive capacity in adult life. The measurement also reflects the pubertal growth spurt, with girls reaching the peak of cerebral growth around 11 years and boys at 14 years [3], as corroborated by Giedd (2008) [4], regarding maximum skull volume.

Currently, head circumference is applied beyond pediatrics in public health, where it serves as a retrospective indicator of early nutritional insults. The evolution towards modern technologies, such as magnetic resonance imaging and tomography, complements traditional anthropometry, offering greater precision in the study of the brain. In adults, this measurement is used in ergonomics (e.g., helmet design) as well as to assess brain reserve; clinically, monitoring head circumference allows for the identification of variations within or outside normal limits, such as microcephaly or megalencephaly, which correlate cranial size to brain mass volume and potential cognitive development. Since a smaller head circumference is correlated with an earlier and more severe manifestation of Alzheimer’s disease in predisposed individuals, cranial anthropometry in mature ages also serves as a risk predictor for cardiovascular diseases, reflecting failures in fetal and infant growth that program metabolism in adult life [3].

### Cranial Capacity, Microcephaly and Megalencephaly

Despite their widespread use, head circumference and cranial capacity (or brain volume) are not equivalent measures, and treating them as interchangeable indicators can lead to interpretation errors, as will be demonstrated throughout this work. The endocranium corresponds to the internal space of the skull, which can be digitally segmented into 3D models to estimate cranial capacity. Brain volume, which reflects the actual size of the brain, is obtained through more complex segmentations, generally using computed tomography or magnetic resonance imaging. Studies indicate that brain volume is, on average, 9.81% smaller than endocranial volume, due to the presence of other tissues, such as meninges and cerebrospinal fluid, in the intracranial space [5].

Microcephaly is a neurological condition characterized by a cranial circumference significantly smaller than the average for the individual’s age and sex, often associated with a reduced brain volume. It can be classified as congenital, present at birth, or post-natal (acquired), developing over time due to factors such as infections or trauma. Epidemiologically, the prevalence of microcephaly is estimated at about 2% of the population when the cranial circumference is two standard deviations below the mean, and 0.1% when it is three standard deviations below [6]. The causes of microcephaly are varied, including genetic mutations, maternal infections during gestation (such as toxoplasmosis, syphilis, varicella-zoster, or rubella), and, in the Brazilian context, the Zika virus, which caused a significant increase in the incidence of microcephaly between 2015 and 2016, with rates ranging from 4% to 8% of live births [6]. The complications associated with microcephaly are serious and can include intellectual disability, developmental delays, epilepsy, cerebral palsy, as well as visual and hearing impairments. In terms of life expectancy, studies indicate that about 82% of individuals with microcephaly do not survive beyond 30 years, with an estimated average life span of 35 years [7]; [8]. Cranial capacity, when measured as brain or endocranial volume, can also be used as a diagnostic criterion, being considered indicative of microcephaly when it is two or three standard deviations below the mean [7]; [9].

Conversely, megalencephaly is characterized by a brain with size and weight significantly above average for the age and sex, usually defined as two standard deviations above the population mean. Clinically relevant cases, however, are often identified when brain volume exceeds three or more standard deviations, as individuals with mild increases (between +2 and +3 SD) may present with normal neurological development. More extreme alterations, such as volumes exceeding 10 standard deviations, are frequently associated with severe conditions, such as mutations in genetic pathways like PI3K-AKT-MTOR, which can result in epilepsy, autism, cortical malformations, and other significant neurological impairments [10].

### Scope and Objectives of the Study

The present study proposed the development and validation of an estimation model for endocranial volume based on a “triple-measure”—the sum of head circumference, sagittal arc, and coronal arc—obtained through Forensic Facial Approximations and supplemented by computed tomography data. The main objective was to demonstrate that the triple-measure offers a significantly more robust correlation with endocranial volume than head circumference alone, serving as a non-invasive, low-cost screening tool for the early identification of atypical cranial–brain volume patterns, such as microcephaly and megalencephaly, across different age groups.

It is important to emphasize that head circumference measurement is already routinely performed in clinical practice, particularly in pediatric follow-up. The proposed approach does not replace existing diagnostic methods but rather complements them by incorporating two additional cranial measurements that can be obtained non-invasively and with minimal effort. Therefore, the model should be interpreted as an experimental screening framework intended to support early detection and clinical decision-making, not as a definitive diagnostic tool. Its purpose is to identify potential deviations from expected cranial–brain volume relationships that may justify further investigation using imaging methods such as computed tomography or magnetic resonance imaging.

## Materials and Methods

### Initial Step - Base Table

#### Initial Case Selection

The initial stage of the present study was based on a set of 100 forensic facial approximations performed between 2013 and 2025, covering a wide variety of cases in adult individuals. This sample includes both sexes and spans multiple geographic and chronological contexts, ranging from the prehistoric era to the 19th century. The inclusion of archaeological and historical remains from various continents provides a comprehensive representation of human cranial morphological variability, establishing a diverse foundation for data analysis, the methodological robustness of which will be addressed in detail further on. These approximations were performed using open-source tools, such as the Blender 3D modeling software (https://www.blender.org/), licensed under the GNU General Public License (GPL) version 2 or higher, ensuring accessibility and transparency in the development and replication of the work. The facial approximations followed two specific approaches over the years: from 2013 to 2021, the work was based on the methodology of Moraes et al. (2014) [11] (Fig. 1, A), and from 2022 to 2025, it followed the steps of Abdullah et al. (2022) [12].

**Figure 1.**
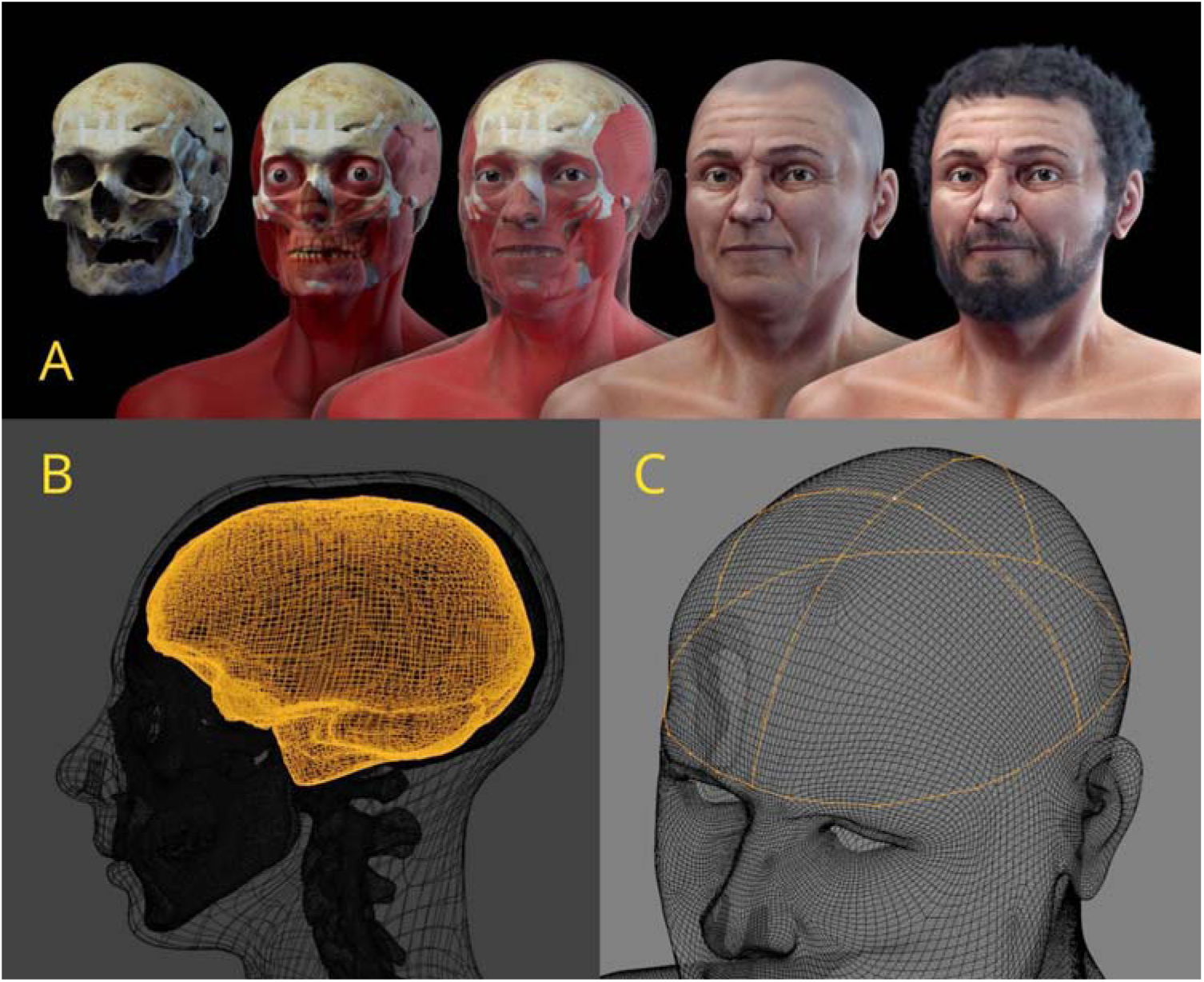
A) Example of forensic facial approximation performed in 2017. B) Endocranial segmentation. C) Triple-measure lines in yellow.

To ensure the consistency and validity of the data analyzed, rigorous inclusion and exclusion criteria were established. The selected facial approximations met the following requirements: clear identification of the individual’s sex and age group, as well as reliable data on the actual scale of the skull, obtained from 3D scans or computed tomography. Initially, facial approximations of very young individuals (pre-pubescent children), or cases where there were doubts about the accuracy of the actual scale of the digital model were excluded. These criteria ensured that the dataset was representative and suitable for the proposed anthropometric analyses, minimizing variables that could compromise the accuracy of the results.

#### Software and Hardware

The development of the study was carried out using open-source tools, which offer flexibility, accessibility, and the possibility of replication across different platforms. The main software employed was Blender 3D (https://www.blender.org/), a robust tool for three-dimensional modeling, animation, and rendering, widely used in areas such as forensic anthropology and computer graphics. To facilitate the importation and manipulation of computed tomography scans, the OrtogOnBlender addon (https://www.ciceromoraes.com.br/doc/pt_br/OrtogOnBlender/index.html) was used, a complement specifically developed to integrate tomography data into Blender version 2.91, a functionality not available in the standard version of the software. This combination allowed for the precise segmentation of anatomical structures, such as the endocranium, and the performance of detailed anthropometric measurements.

The collected data were organized and stored in tables in CSV (Comma Separated Values) format, a simple text format that facilitates interoperability and data sharing. For generating graphs and statistical analyses, the Python programming language (https://www.python.org/) was used, which offers powerful libraries such as Matplotlib and Pandas for data visualization and manipulation. The choice of free software, such as Blender and Python, reflects the study’s commitment to accessibility and reproducibility, allowing the methods to be replicated on popular operating systems, such as Linux Ubuntu (used in this work), Windows, and macOS.

The hardware used was a Lenovo Ideapad S145, equipped with an Intel Core i7-8565U processor, 20 GB of RAM, and an NVIDIA GeForce MX110 graphics card. This configuration was sufficient to support the computational demands of 3D model and tomography processing, ensuring efficiency in task execution.

#### Endocranial Volume Estimation

The segmentation of the endocranium, essential for estimating cranial capacity, was performed differently depending on the origin of the data. In cases where the forensic facial approximation was based on computed tomography scans, the endocranium was segmented directly from the internal negative of the skull’s 3D mesh, extracted from the tomography (Figure 1, B). The resulting structure was closed using Blender’s Remesh tool, which allows the reconstruction of 3D meshes to ensure a continuous surface suitable for volumetric calculations (https://docs.blender.org/manual/en/latest/modeling/modifiers/generate/remesh.html). The endocranial volume was then calculated with the volume measurement tool of the 3D Print Toolbox addon (https://docs.blender.org/manual/en/3.6/addons/mesh/3d_print_toolbox.html), which provides precise results in cubic centimeters.

Endocranial volume was considered the primary measurement variable throughout the study. When brain volume values are presented, they represent derived estimates obtained from endocranial measurements using the standardized reduction factor described below, allowing comparison with studies that report brain tissue volume instead of endocranial capacity.

The CT datasets used in this study originated from standard clinical acquisitions with typical matrix resolution of 512 × 512 pixels. Because the scans were obtained from heterogeneous medical environments, specific acquisition parameters such as scanner model, voltage, exposure time, and slice thickness varied across cases. However, these variations do not significantly affect the estimation of global endocranial volume, which depends primarily on geometric segmentation of the cranial cavity rather than on fine tissue contrast. The use of heterogeneous clinical datasets reflects real-world conditions and does not compromise the volumetric measurements obtained. Brain volume estimates were derived from endocranial volume using a reduction factor of 9.81%, based on previously published empirical observations relating endocranial cavity volume to brain tissue volume [5]. As will be shown in two sex-specific meta-analyses presented below, even when accounting for the differences between endocranial and brain volumes, the brain is approximately 10% smaller than the endocranium, consistent with [5]. For further details, see Tables S6 and S7 in the Supplementary Material.

It is important to note that endocranial volume constitutes the primary variable used in the proposed model (Tables 2 and S8). Brain volume values presented in Table 1 were derived solely for comparative purposes with heterogeneous literature reports, which include both cranial capacity and brain measurements. Therefore, the use of different terms reflects the distinct roles of these datasets rather than a conceptual inconsistency.

**Table 1:** Comparison between literature-derived brain volume equivalents (meta-analysis) and the present sample.

| Measure | Meta-Analysis | Present Study |
| --- | --- | --- |
| Head Circum. (Male) | 56.34 cm (± 2.33) | 56.40 cm (± 2.14) |
| Head Circum. (Fem.) | 54.49 cm (± 2.17) | 54.57 cm (± 2.27) |
| Brain (Male) | 1238 cm<sup>3</sup> (± 103) | 1282 cm<sup>3</sup> (± 134) |
| Brain (Fem.) | 1119 cm<sup>3</sup> (± 93) | 1188 cm<sup>3</sup> (± 111) |

For externally digitized skulls only (without tomography), the anatomical deformation technique [13] was employed. In this method, a virtual donor skull, obtained from a computed tomography (CT) scan, was deformed to structurally align with the target skull of the facial reconstruction. After deformation, the internal negative of the adjusted skull was segmented, following the same steps described for CT-based cases, ensuring consistency in volumetric calculations. This approach allowed for the acquisition of precise data even in cases where CT scans were unavailable, expanding the method’s applicability.

Surface digitizations were obtained using photogrammetry following the protocol previously described in Moraes et al. [11], which has been applied in clinical and forensic contexts, as further addressed in the Discussion section. This approach provides sufficient geometric accuracy for external cranial morphometric measurements such as those employed in the present study.

For these cases, multiple virtual donor skulls derived from computed tomography datasets were used. Donor selection was performed according to the sex and ancestry of the target individual whenever such information was available, providing anatomically consistent reference structures for the deformation process. Because the anatomical deformation is guided by the external cranial morphology and measurements of each case, the final adjusted geometry aligns with the target skull rather than retaining donor-specific characteristics, thereby minimizing potential representational bias.

#### Measurement of Head Circumference and Sagittal and Coronal Arcs

The head circumference measurement was taken in the region immediately above the eyes and ears, according to established standards in the literature [14]. In addition, the sagittal and coronal arches were measured, respecting the lower limits of each arch and the center of their respective axes (Figure 1, C). These measurements were extracted using the Measureit tool from Blender (https://docs.blender.org/manual/en/3.0/addons/3d_view/measureit.html), which allows precise measurements directly in the 3D environment, with results exported to CSV tables. This approach guaranteed the standardization of the measurements and facilitated comparison with reference data from population studies.

Linear regression models were explored to evaluate relationships between cranial measurements and endocranial volume and to derive preliminary estimation formulas. These models were used as intermediate analytical steps to integrate heterogeneous datasets and to support the development of the final lookup table and graphical reference system presented in the Results section. The resulting formulas were not intended as standalone predictive tools but as components of a broader empirical modeling process whose clinical application is limited to screening purposes rather than definitive diagnosis.

Since the protocol for measuring head circumference has already been established by Macchiaverni & Barros Filho (1998) [4], the sagittal and coronal arcs must be obtained using the circumference line as the basal reference for the lower limits of each measurement.

##### Sagittal Arch

The measurement begins at the glabella, follows the sagittal midline over the vertex (the highest point of the skull), and ends at the external occipital prominence, at the lower limit of the previously measured circumference. The tape measure must maintain direct contact with the scalp, following the anatomical curvature of the skull.

##### Coronal Arch

The tape should be positioned transversely, starting at the region above the ear (temporozygomatic joint), passing over the vertex, and ending at the homologous region on the opposite side. This measurement represents the largest transverse arc between the lateral limits of the head circumference.

##### Practical Procedure

An efficient way to ensure precision is to mark the intersection points on the tape measure during the circumference measurement. The specialist may use a non-toxic pigment to mark these points or, alternatively, tie an auxiliary string along the circumference line, using it as a fixed delimiter for the start and end of the sagittal and coronal arc measurements.

#### Meta-Analyses of Head Circumference and Cranial Capacity

The literature-derived datasets described in this section were used as external reference parameters for model contextualization rather than as part of a formal systematic meta-analysis. The studies included were identified through targeted searches in Google Scholar and PubMed using terms related to head circumference, cranial capacity, and brain volume in adult populations. To contextualize and complement the obtained data, two comprehensive meta-analyses were performed. The first meta-analysis focused on head circumference and included nine studies [15]; [16]; [17]; [18]; [19]; [20]; [21]; [22]; [23], totaling a sample of 2,348 male and 2,074 female individuals. This analysis provided a robust basis for establishing parameters of normality and assessing deviations in specific cases.

A second meta-analysis focused on cranial capacity, compiling ten studies [24]; [25]; [4]; [26]; [27]; [28]; [29], with a total sample of 3,496 male individuals and 3,410 female individuals. This analysis allowed for the comparison of endocranial volume data obtained in the present study with established reference values in the literature, reinforcing the validity of the measurements and highlighting the importance of considering cranial capacity as a complementary indicator to head circumference.

Because the reference studies reported heterogeneous variables (including endocranial volume, cranial capacity, and brain volume), values were standardized when necessary to ensure comparability. These literature-derived datasets were used exclusively for contextual validation and compatibility assessment of the adult sample, rather than as direct inputs for model construction.

#### DICOM Data Selection and Preparation

For a preliminary validation pilot test, computed tomography datasets from 10 individuals were randomly selected from a bank of anonymized scans, ensuring diversity of ancestries and representativeness. Each dataset consisted of a full series of DICOM (Digital Imaging and Communications in Medicine) slices corresponding to a complete cranial examination. The segmentation of the endocranium and subsequent measurements followed the procedures previously described, using the OrtogOnBlender addon to import and process the tomography data in Blender. This process involved converting the DICOM images into 3D models, segmenting the endocranium, and applying measurement tools to obtain precise anthropometric data.

### Complementary Step - Scope Expansion for Pediatric and Neonatal Use

Although the facial approximation sample comprised a number of 100 individuals, the remaining samples related to children and babies were significantly smaller, which makes it impossible to use them for establishing a statistically robust graph. To overcome this data deficiency, studies were used that contained information on head circumference vs brain volume/cranial capacity vs age [30]; [31], and studies with cranial capacity vs age data [32]; [33]; [34]. These studies were identified through targeted searches in Google Scholar and PubMed using terms related to brain volume, cranial capacity, and head circumference across developmental stages. Data were extracted from tables when available and, when necessary, digitally approximated from graphical representations when numerical tables were not available published in the original articles. Due to the heterogeneity of the pediatric datasets and the absence of individual-level measurements fully compatible with the proposed model, representative age-group subsets described in the source studies were selected, and approximate mean values were derived to position these groups along the theoretical curve. These values were then integrated with the proportional relationships derived from the sagittal and coronal arc measurements proposed in the present study, allowing the construction of the reference regions shown in Figure S8.

### Final Stage - Presentation

For the creation of the analog lookup table, the Inkscape software was used to create a PDF file for later printing. Regarding the development of the online consultation page, which in practice functions as an app; an HTML code (https://www.w3.org/TR/html) with JavaScript (https://developer.mozilla.org/en-US/docs/Web/JavaScript) was used for the calculations and the ChartJs package for the graphical presentation (https://www.chartjs.org/).

### Methodological Scope and Intended Use

The proposed model is explicitly descriptive and intended as a screening tool rather than an inferential framework. Its purpose is to identify deviations from expected cranial–brain volume patterns that may warrant further investigation, not to estimate population parameters or establish causal relationships.

The measurements required by the model are based on procedures already used in clinical contexts (e.g., head circumference), with the addition of two complementary cranial arcs, reinforcing its intended role as a practical adjunct rather than a standalone diagnostic method.

Future studies may explore alternative non-parametric or distribution-sensitive approaches, such as quantile regression or locally weighted regression (LOESS), to further refine boundary behavior in regions of low sample density. However, the present approach was intentionally designed to prioritize interpretability, accessibility, and practical screening utility over inferential complexity.

## Results

### Cranial Capacity and Triple-Measure

The present study performed anthropometric measurements on 100 forensic facial approximations, comprising 65 men (Table S1) and 35 women (Table S2). In addition to head circumference and cranial capacity (measured as brain and endocranial volume), sagittal and coronal arc measurements, as well as the triple measure (sum of head circumference, sagittal arc, and coronal arc), were collected.

Regarding head circumference, the final value was 56.40 cm (± 2.14) for male individuals and 54.57 cm (± 2.27) for female individuals, values very close to the meta-analysis. Cranial (brain) capacity resulted in 1282 cm³ (± 134) for males and 1188 cm³ (± 111) for females, also consistent with meta-analyses. The triple measure (Head Circumference + Sagittal Arc + Coronal Arc) resulted in 110.27 cm (± 4.56) for males and 107.28 cm (± 3.35) for females. The overall averages were also provided (Table S3).

### Head Circumference Meta-Analysis

The meta-analysis of head circumference for male individuals covered nine studies, totaling a sample of 2348 individuals. These studies, conducted between 1991 and 2024, provided consistent data on cranial circumference in diverse populations. The resulting weighted average was 56.34 cm (± 2.33), indicating a range of variation that reflects the anatomical diversity among the analyzed individuals. This average serves as a reference parameter for evaluating deviations in specific cases, such as those associated with neurological conditions, and was used to contextualize the results of the present study (Table S4).

For the female sex, the meta-analysis included eight studies, with a total of 2074 individuals. The weighted average head circumference was 54.49 cm (± 2.17), a value slightly lower than that observed for men, consistent with anatomical differences between the sexes. This analysis reinforces the importance of considering gender specificities when evaluating anthropometric measurements, especially in clinical and forensic contexts, where the identification of deviations from expected patterns, such as microcephaly or megalencephaly, may depend on comparisons with population standards (Table S5).

### Meta-analysis of Cranial Capacity

The meta-analysis of cranial capacity for male individuals involved ten studies, totaling 3496 subjects. This analysis differentiated between brain volume and endocranial volume measurements, recognizing that brain volume is, on average, 9.81% lower than endocranial volume due to the presence of non-cerebral tissues in the intracranial space. The weighted average for brain volume was 1238 cm³ (± 103), a value that reflects the expected variability in adult male populations and serves as a basis for comparisons with individual cases, such as those analyzed in the present study (Table S6).

For the female sex, the meta-analysis of cranial capacity also involved ten studies, with a total sample of 3410 individuals. The weighted average of brain volume was 1119 cm³ (± 93), a value lower than that of men, consistent with anatomical gender differences. This analysis highlighted the importance of using gender-specific data to assess cranial capacity, especially in contexts where clinically relevant deviations may be present, requiring precise comparisons with population standards (Table S7).

### Comparison of Results: Meta-Analyses versus Present Study

The compatibility of the present study with meta-analyses highlights a good distribution of the sample (Table 1). Therefore, it can be used as a basis for the study of the compatibility between the triple-measure and endocranial volume.

### Composition of the Triple-Measure Base Table

#### Regression and Simple Formulas

Initially, 6 distribution options were plotted to evaluate the R and R² of each (Figure S3). Head circumference, as it is popular, and double and triple measures, as they are more robust, received a simple linear regression formula (Y=mX+b) for calculating the potential volume of the endocranium based on them.

- The comparison between head circumference versus the endocranium generated R = 0.76 and R² = 0.57, indicating a strong positive correlation.

○ Calculation formula: Endocranium = (47.61 × HeadCircumference) − 1271.91
- ● The comparison between the sagittal arc and the endocranium resulted in R = 0.68 and R² = 0.45, indicating a strong positive correlation.
- ● The comparison between the coronal arc and the endocranium resulted in R = 0.52 and R² = 0.27, indicating a moderate positive correlation.
- ● The comparison between the joint measure of head circumference + sagittal arc versus the endocranium resulted in R = 0.83 and R² = 0.70, indicating a very strong positive correlation.

○ Calculation formula: Endocranium ≈ 17.452 × (HeadCircumference+SArc) − 388.33
- The comparison between the joint measure of head circumference + coronal arc versus the endocranium resulted in R = 0.82 and R² = 0.68, indicating a very strong positive correlation.

○ Calculation formula: Endocranium ≈ 17.921 × (HeadCircumference+CArc) − 416.12
- The comparison between the joint measure of head circumference + sagittal arc + coronal arc versus the endocranium resulted in R = 0.83 and R² = 0.69, indicating a very strong positive correlation.

○ Calculation formula: Endocranium ≈ 21.322 × (HeadCircumference+SArc+CArc) − 992.36

In 98% of the cases, the sagittal arc was larger than the coronal arc, demonstrating a consistent pattern potentially exploitable by specialists (Figure S4).

Although the results of just two measures—in this case, head circumference + sagittal arc—already showed a very strong correlation, including slightly above the triple measure (head circumference + sagittal arc + coronal arc), the pattern of difference between the arcs, coupled with the safety of generating an extra measure for confrontation, led to the triple measure being chosen for a more detailed approach.

The F-test for comparing the R² values of the endocranium versus head circumference and the endocranium versus the triple measure was calculated using the formula (Figure S5):

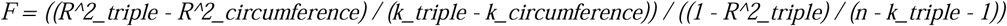

For a significance level of α = 0.05, the critical F value (from F tables) is approximately 3.09. Since F ≈ 17.03 is much greater than 3.09, the difference is statistically significant (p < 0.001), indicating that the triple measurement model is significantly better at explaining endocranial volume.

The problem with formulas based on simple linear regressions is that, although the standard deviations work well in denser regions of the sample, in less dense regions, with less expected variation, such as cases with potential microcephaly, the standard deviation tends to be exacerbated, not reflecting such variation (Figure S6, A). One way to solve part of the problem is by creating a moving average that reflects the characteristics of the sample region, but although this partially resolves the issue of variation, the part related to microcephaly tends to be neglected, in addition to exacerbated peaks in the standard deviations due to the absence of attenuation (Figure S6, B). A solution that satisfies the regional characteristics of the sample and its amplitude, while respecting the standard deviations, is the correction of the lines by attenuation like a bézier curve. The final graph, with the simplified curve and containing two standard deviations, which cover ∼95% of the sample, could be configured (Figure S6, C).

To facilitate the analysis, a table was created with the sample values, allowing consultation of the potential endocranial volume based on the sum of the head circumference, sagittal arc, and coronal arc (triple-measure), including their respective standard deviations (Table S8). The objective of the table is to provide a simplified consultation system based on the triple-measure, with rounded results. An example could be an individual whose measurements resulted in: 55.73 cm for head circumference, 29.29 cm for the sagittal arc, and 26.68 cm for the coronal arc, totaling 111.70 cm. Rounding the value reaches 112, or an average of 1443 cm³, with the respective standard deviations. Following this, comparisons of the projections with the actual volume measurements will be discussed.

#### Previous Validation of the Triple-Measure Table (Adults)

For the data validation test, 10 computed tomography scans of adults were used, randomly selected from an anonymized database. The DICOM files were reconstructed, including bone and soft tissue factors, with the endocraniums segmented and the triple measurement performed. The data were then organized and compared with the general table (Table S8).

The compatibility of the actual value with the projection for the triple measurement was assessed. In Case 1, the endocranium measurement resulted in 1253 cm³, and the triple measurement was 105.11, rounded to 105. In the table, the value was slightly higher, 1265 cm³, but very close to the actual value, being colored green because it was closer to the mean than to the -1 SD limit. In Case 2, the result was 1267 cm³ with a triple measurement of 101.57, rounded to 102. The actual volume was between +1 SD (1232) and +2 SD (1309); however, as the average of the two is 1270.5 cm³ and the actual volume is below this value, it is closer to +1 SD and was colored yellow. The same logic was applied to the remaining cases.

A graph was plotted based on the standard deviation of each case (Figure 2), allowing visual confirmation of the limits of each standard deviation and the rationale for the chosen coloration. Of the 10 cases, 5 were closer to the mean, 4 were closer to ±1 SD, and 1 was close to ±2 SD. When considering only the standard deviations, 7 cases fell between 0 and ±1 SD, and 3 fell between +1 SD and +2 SD, without exceeding +2 SD; thus, all remained within the table/curve (Table S8, Figure S6, C). This demonstrates the robustness of the approach in relation to the pilot sample used, which consisted of living and real individuals, rather than forensic facial approximations.

**Figure 2:**
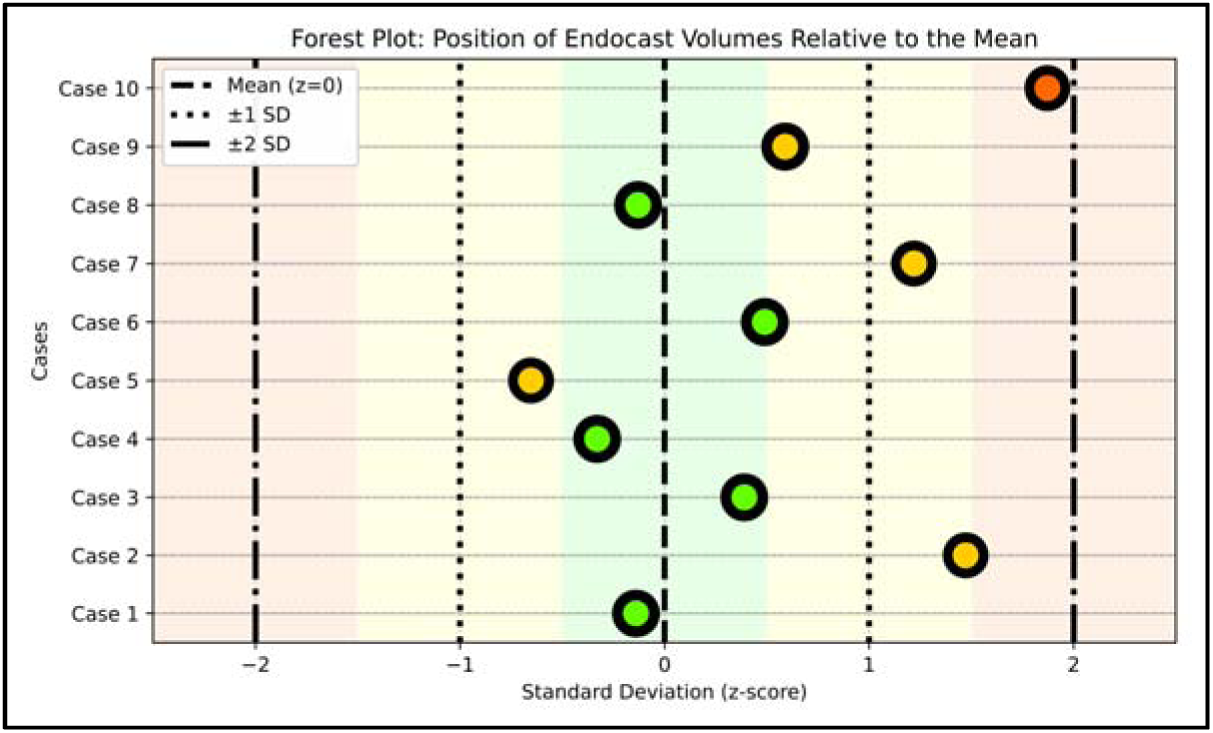
Distribution of the actual measurements in relation to the mean and standard deviations of Table S8. Point colors indicate proximity to reference limits: green = closest to the mean; yellow = closest to ±1 SD; red = closest to ±2 SD.

Another graph was plotted (Figure S7), using the actual values as a basis (dotted black line) comparing it with the projections derived from the prediction based on the head circumference formula (in red), the triple-measure formula (in green), and the simplified triple-measure table (in blue). Of the 10 skulls evaluated, 9 were within two standard deviations (for general cranial capacity), meaning they can be considered within the region of normality; only one of them exceeded this value. The comparison evidences the greater compatibility of the triple measure in relation to the projection made from the isolated head circumference, with a small advantage of the table over the formula derived from simple linear regression.

The original sample contains a classic case of microcephaly and some other suspicious cases, including that of an adult male skull, with characteristics that highlight the limitations of analysis by head circumference, as at 53.5 cm, it could be considered within the normal range for males, as it is in the -1.22 SD region. As it is a suspicious case, at the limit of the literature, which would be -2 SD, it becomes a well-selected evaluation to test the approaches available in the present work. The cranial capacity was raised to 1101 cm³ for the endocranium, which converted into cerebral volume resulted in 993 cm³, which could already be indicated as a potential case just for being a mature adult and being below 1000 cm³ [7]. The other measurements resulted in 24.62 cm for the sagittal arc (-2.21 SD), 22.51 cm for the coronal arc (-1.94 SD), totaling the triple measure at 100.63 cm (-2.11 SD). A comparison with the formulas and the table clearly illustrates the limitations of the calculation based on head circumference (see Table S10), which indicated a volume of 1275 cm³ for the endocranium, placing it within a negative standard deviation, both in the meta-analysis and for the most robust and highly academic study of the analysis [8] (Table 9). The formula of head circumference + coronal arc was the one that most closely approximated the actual result, indicating 990 cm³ and comfortably below two negative standard deviations. The triple table showed the position in a borderline region, at -1.93 SD, mainly due to the simplification of the calculation (triple measure of 100.63 was rounded to 101), because by the system’s standard deviation, calculated precisely, it would yield -2.11 SD, which proved consistent with the study by [28], which resulted in -2.14 SD (Table S10). However, in a screening case, the simple measure would already be sufficient for more careful observation, additional calculations, and a potential request for computed tomography for a deeper analysis. Furthermore, the possibility of calculating the volume with different combinations of measures allows specialists to estimate the structure even in the absence of one of the three essential measures, as in the case of data collection failure.

#### Additional Statistical Validation of Predictive Performance

To further evaluate the predictive performance of the proposed approach, additional statistical analyses were conducted using the same independent set of 10 computed tomography cases described above. Predicted endocranial volumes were extracted from Table S8, which was generated exclusively from the 100 forensic facial approximations, ensuring independence between calibration and validation datasets.

Agreement between predicted and observed values was assessed using Pearson correlation, intraclass correlation coefficient (ICC; two-way mixed model, absolute agreement), Bland–Altman analysis, mean absolute error (MAE), and root mean square error (RMSE). These metrics were also calculated for predictions based solely on head circumference to allow direct comparison between methods.

The triple-measure model demonstrated strong agreement with observed endocranial volumes (ICC = 0.84), substantially higher than head circumference alone (ICC = 0.57). Pearson correlation was also higher for the triple measure (r = 0.87, p = 0.001) compared to head circumference (r = 0.81, p = 0.004). Bland–Altman analysis revealed lower mean bias for the triple measure (41.5 cm³) compared with head circumference (96.2 cm³), with narrower limits of agreement (−103 to +186 cm³ vs. −83 to +275 cm³). Error metrics similarly favored the proposed method (MAE = 65 cm³; RMSE = 81 cm³) relative to head circumference alone (MAE = 96 cm³; RMSE = 129 cm³).

The positive bias observed in both methods is consistent with the known limitation associated with measurements obtained from digital cranial models without hair volume, as discussed later in the manuscript. Overall, these results indicate that the triple-measure approach provides more accurate volumetric estimates than head circumference alone, supporting its potential utility as a complementary screening-oriented tool.

The computational implementation of the statistical validation procedures is provided in Supplementary Code S1.

#### Expansion for Pediatric Use

During the pilot study for observing the tool and rereading some references, it became evident that there was a possibility to expand the scope of the tests, since, when the baby reaches 2-4 weeks of age, their brain already represents more than a third of the final size, or 36%; in the first year of life it increases to 72%, at 2 years to 83% [33]. The growth rate remains significant, reaching 95% of the total volume at 6 years of age [32] and peaking, that is, 100% of the volume at ∼11 years in girls and at ∼14 years in boys [4].

The studies together totaled aggregated datasets comprising several hundred individuals across developmental stages, so it was necessary to integrate heterogeneous datasets, based on projections and on brain/cranial volume. To overcome this lack of data, studies containing information on head circumference versus brain volume/cranial capacity versus age [30]; [31], as well as studies with data on cranial capacity versus age [32]; [33]; [34], were used. The graphical data referenced from these studies were not reproduced in the present work due to licensing restrictions; however, all corresponding plots and datasets can be directly consulted in the original publications cited above. Initially, a projection of the triple measure (TM) was performed based on head circumference. According to the overall average found in the present study (see Table 5), the average adult endocranial volume is 1385 cm³ and the head circumference is 55.8 cm. The average sagittal arc is 28.08 cm, or 50.32% of the head circumference (HC); and the coronal arc is 25.32, or 45.38% of the head circumference, therefore the formula for projecting the triple measure would be the following:

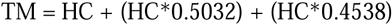

This projection assumes that proportional relationships between head circumference and cranial arcs observed in adults provide a reasonable approximation for developmental stages.

Using this formulation, TM values were estimated for datasets in which only head circumference was available. In cases where cranial capacity or brain volume and age were directly reported in the literature, these values were plotted without transformation, assigning TM to the horizontal axis and volume to the vertical axis. Age information, when available, was represented as vertical reference lines to preserve developmental context across the integrated dataset.

The data projected from the triple measure projection using head circumference showed a good distribution, reducing the starting point of the triple measure which was previously 96, to 50 cm (Figure S8).

#### Prior Validation of the Triple-Measure Model (Neonates and Children)

Just as a pilot test was established in the adult age group to test the curve’s accuracy, the same was implemented for the neonate and infant region. Facial approximations of children (n=3) from ∼6 to ∼14 years of age and infants (n=4), with no established age, were chosen, as they are archaeological skulls. Upon distributing the measured individuals on the plotted graph, almost all approached the expected mean (Figure 3).

**Figure 3:**
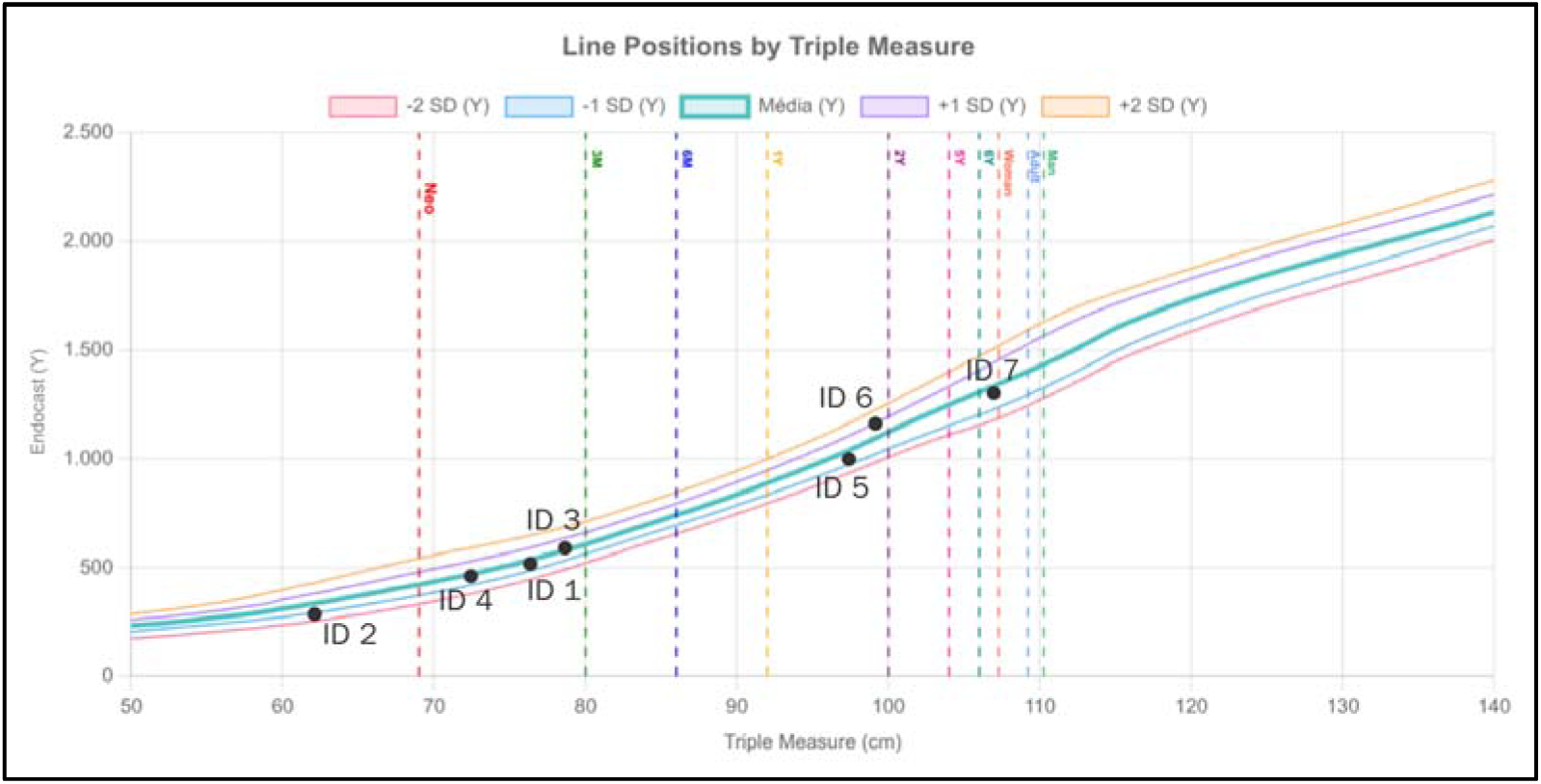
Empirical anatomical data and projected relationship between triple-measure values and cranial capacity across developmental stages. Data points include measurements derived from the present study and literature-based reference groups, while the continuous curve represents the modeled projection generated from the proposed framework.

### Data and Tools Presentation

#### Final Triple-Measure Table

Since the pilot measure data from the 10 blind cases showed a steeper curve trend in skulls with larger volume, a small correction was made to conform the data to the expected volume and the final limit of the triple-measure increased from 121 to 140 cm.

In light of this adjustment, the adult study, which was limited from 922 cm³ to 1840 cm³, comprising neonates, children, and individuals with significant megalencephaly, increased the range, currently from 175 cm³ to 2280 cm³ (Table 2). The expected means and standard deviations were also established for neonates, 3-month-old infants, 6-month-old infants, 1-year-olds, and children aged 2 years, 5 years, and 6 years (Table 3).

**Table 2:**
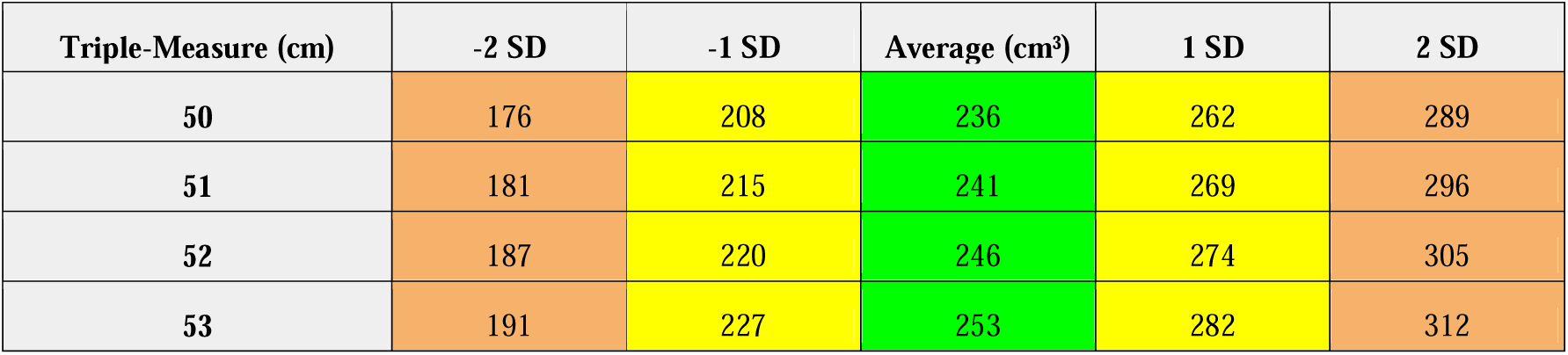
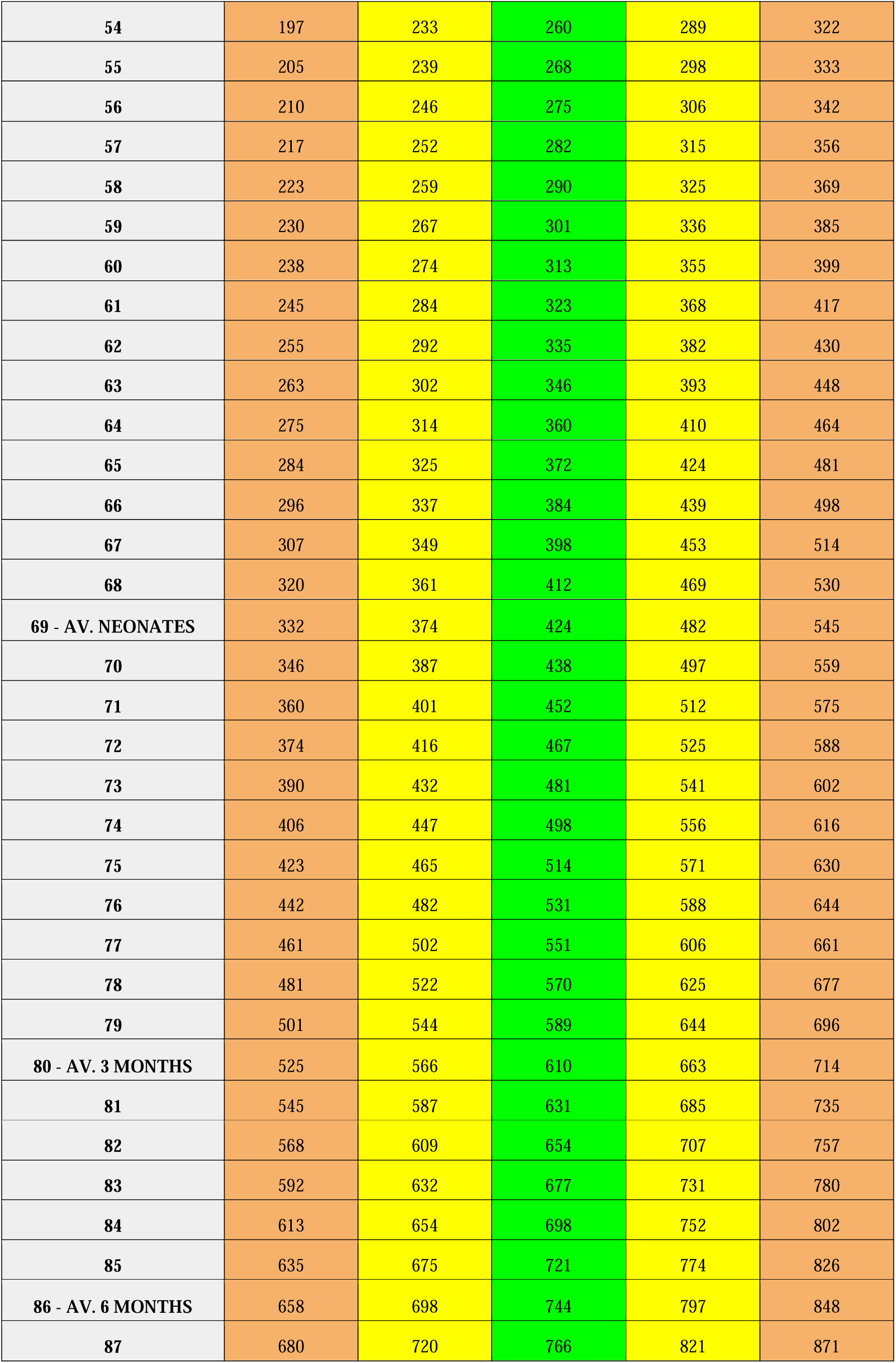
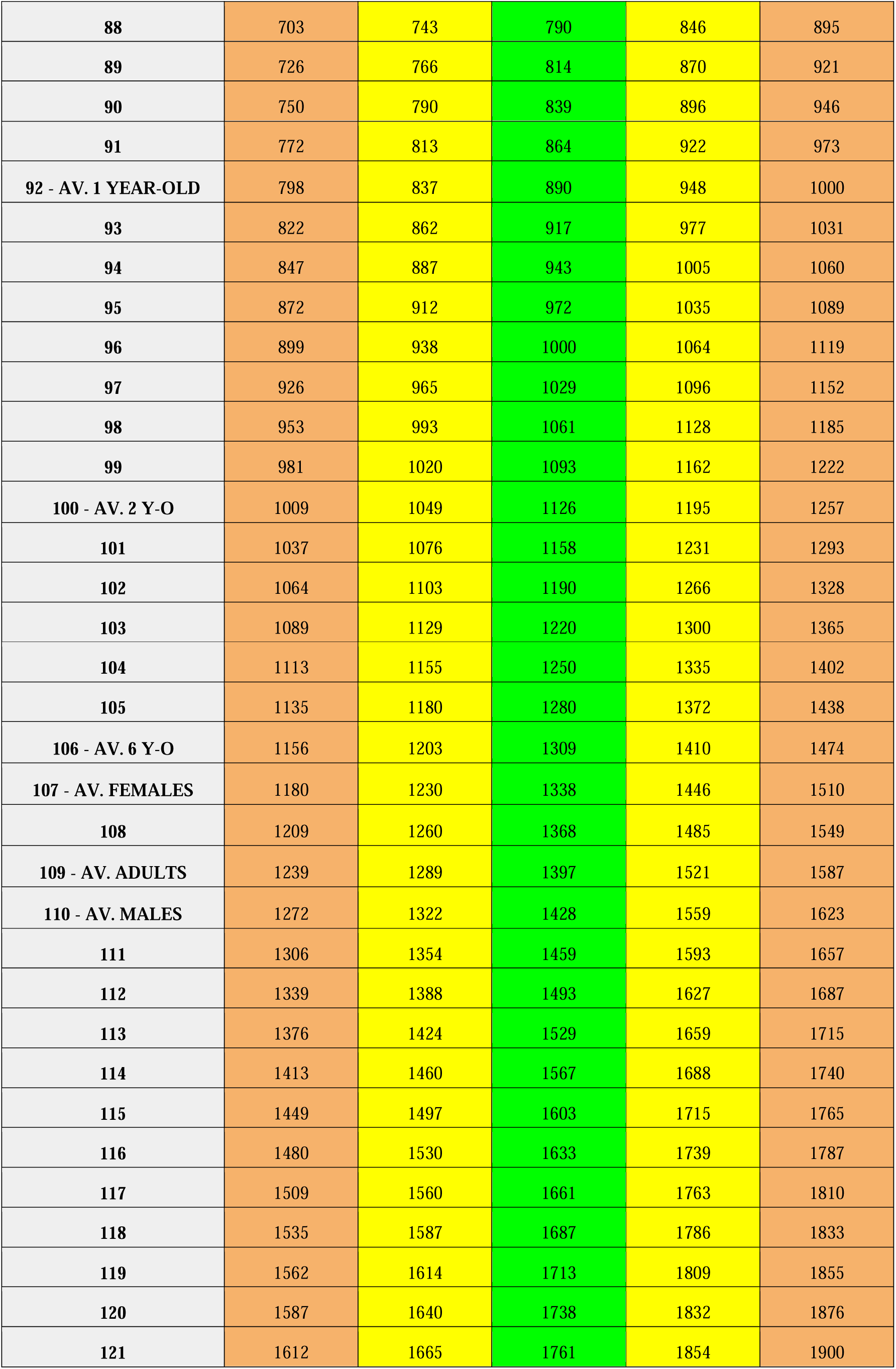
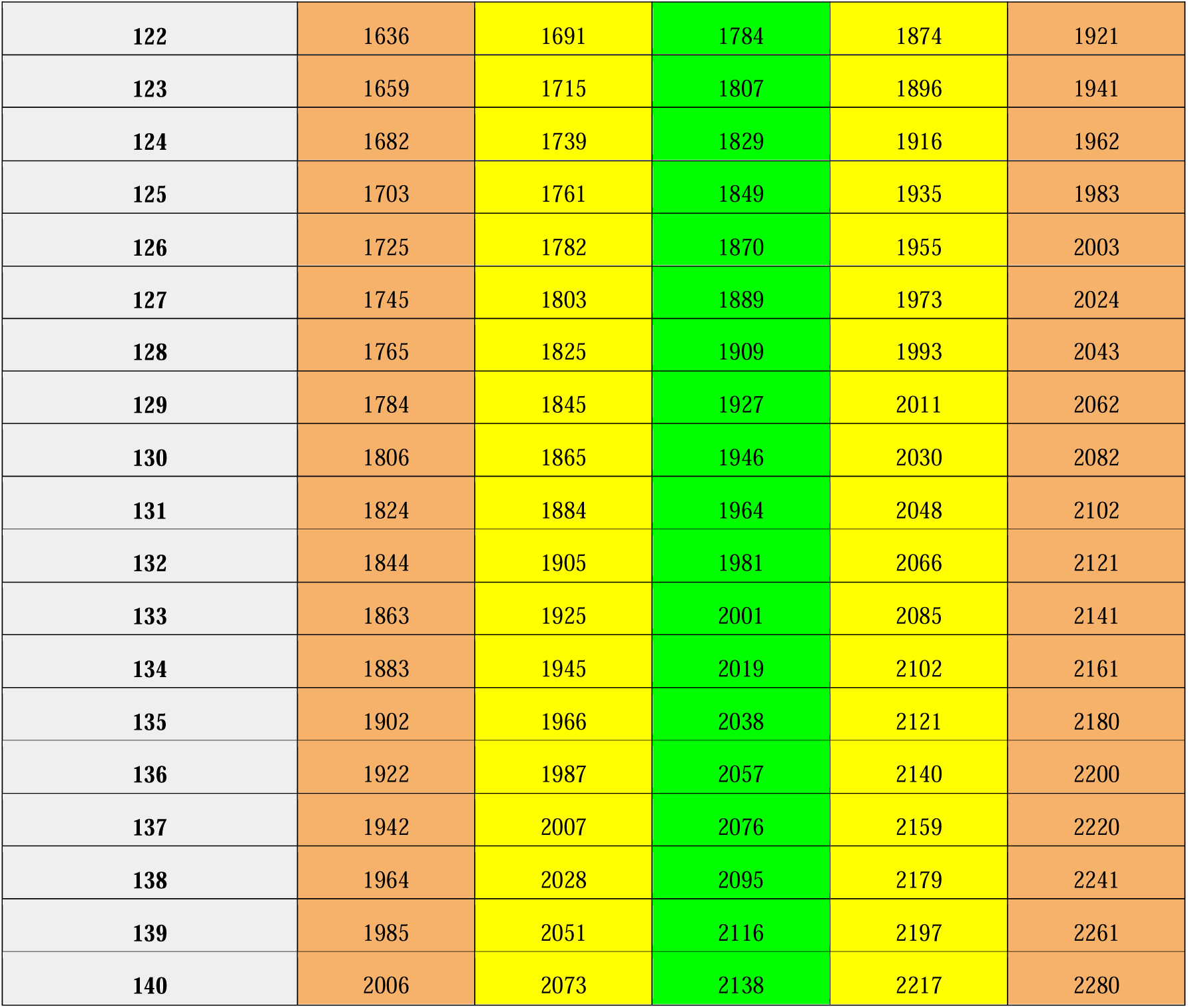
Triple measure and corresponding endocranial volume estimates across neonates, children, and adults.

**Table 3:** Triple measure corresponding to each marker, with the standard deviation.

| Marker | Mean Triple Measure (cm) | Standard Deviation (cm) |
| --- | --- | --- |
| Neonates | 69.00 | ± 4.00 |
| 3 Months | 80.00 | ± 3.50 |
| 6 Months | 86.00 | ± 3.00 |
| 1 Year | 92.00 | ± 3.00 |
| 2 Years | 100.00 | ± 3.00 |
| 5 Years | 104.00 | ± 2.00 |
| 6 Years | 106.00 | $\pm 2.00$ |
| Females | 107.28 | $\pm 3.35$ |
| Adults | 109.22 | $\pm 4.40$ |
| Males | 110.27 | $\pm 4.56$ |

#### Calculation Page

Although the triple-measure/volume table forces an analog and simple way to consult the values, the authors sought to facilitate the process even further, composing an html page with javascript that allows the data to be filled in, the calculation, and the visualization of the patient on the two-dimensional graph.

The html can be accessed at https://script.google.com/macros/s/AKfycbxyTSaETS30sMLL8gCbrYFei5AqT2jVbkzPKt6Cem04Ulj4zETwaI3BLpAJGEm14MT/exec and used for the proposed calculation (Figure 4). If the user is interested, it can be downloaded, saving the page as a file and used from a computer or even a smartphone, practically functioning as a cross-platform offline app, namely Windows, MacOS, Linux, Android, IOS, and others with html and javascript support.

**Figure 4:**
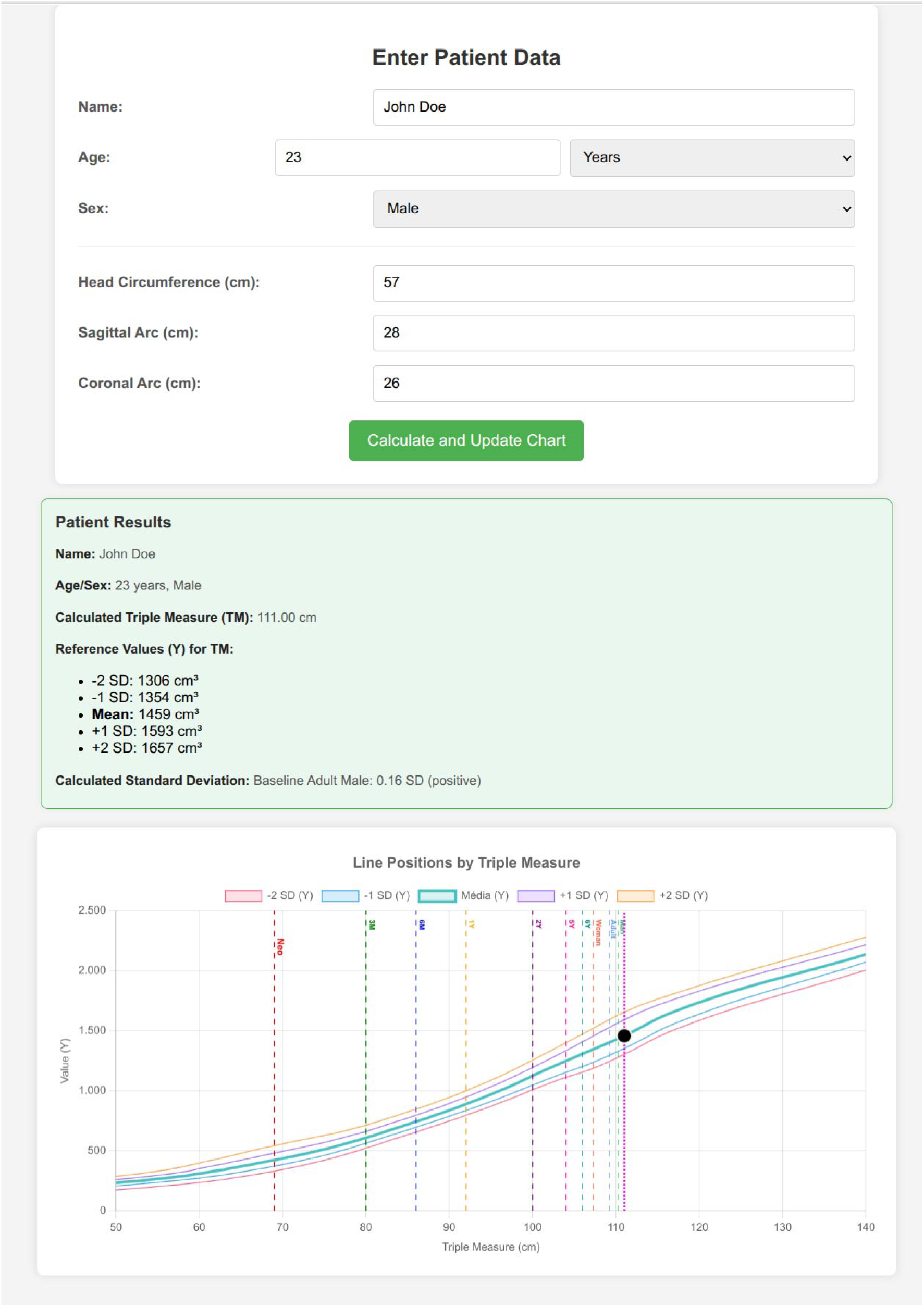
Graphical interface for the triple-measure calculation.

As illustrated in Figure 4, there is an interface for filling in patient data. In the “Name” field, which is optional, the user fills in the name of the person being evaluated. In “Age,” there is the option of years or months; as this is an adult, the choice was obviously “Years.” Sex is also optional, but in this case, “Male” was selected because it refines the results, given that statistically, the adult male skull is slightly larger than the female one. Next, the data for the triple-measure are filled in, which are the head circumference, the sagittal arc, and the coronal arc. Once all the information is satisfied, the user can click on “Calculate and Update Chart.”

As soon as the button is pressed, a frame with the data collected from the patient is presented to the user, such as the triple measure, calculated as 111.00 cm, and the average expected cranial capacity, as 1459 cm³ with the respective standard deviations. Below is presented the patient’s standard deviation in relation to the belonging group, in this case, resulting in +0.16 SD. The curve graph is updated with a black circle corresponding to the patient. The tool issues an alert, both in case of potential microcephaly and potential megalencephaly.

Since this tool is still under development and was designed for screening rather than diagnostic purposes, an ‘Important Notice’ text has been placed at the bottom of the calculator to highlight these characteristics and limitations:

#### Important Notice: Tool Under Development

This site is a tool under development, intended exclusively for growth monitoring and initial screening of microcephaly and megalencephaly cases.

WARNING:

The data and results presented here DO NOT constitute a medical diagnosis.

This tool DOES NOT replace evaluation and clinical judgment by a qualified healthcare professional. Any health-related decision or assessment must be performed by a specialist physician.

The developer of this tool assumes no responsibility for any interpretations, decisions, or consequences resulting from use of the data generated here without proper validation and professional medical follow-up.

## Discussion

The present study addressed the development of the triple measure, composed of the sum of head circumference, sagittal arc, and coronal arc, obtained through forensic facial approximation, as an innovative tool for estimating endocranial volume and assisting in the screening of conditions such as microcephaly and megalencephaly. The results demonstrated that the triple measure exhibits a very strong correlation with endocranial volume (R = 0.83, R² = 0.69), significantly surpassing isolated head circumference (R = 0.76, R² = 0.57), as confirmed by the F-test (F ≈ 17.03, p < 0.001).

Additional validation analyses performed on an independent set of computed tomography cases further supported the predictive performance of the model. Agreement between predicted and measured endocranial volumes was high (ICC = 0.84), substantially exceeding that obtained using head circumference alone (ICC = 0.57). The triple measure also demonstrated lower prediction errors (MAE = 65 cm³; RMSE = 81 cm³) compared with head circumference (MAE = 96 cm³; RMSE = 129 cm³), as well as reduced bias in Bland–Altman analysis. These findings indicate improved accuracy and consistency of the proposed approach while maintaining methodological simplicity.

This superiority was particularly notable in borderline situations, such as an adult male skull with signs consistent with microcephaly, where head circumference (53.5 cm, −1.22 SD) suggested normality, but the triple measure (100.63 cm, −2.11 SD) and the actual endocranial volume (1101 cm³, equivalent to 993 cm³ of brain volume) indicated a potential clinically relevant deviation, aligned with diagnostic criteria. This finding reinforces the sensitivity of the triple measure for detecting subtle cranial–brain deviations that may go unnoticed by traditional methods, fulfilling the screening objective by signaling the need for additional exams, such as computed tomography.

The robustness of the sample of 100 forensic facial approximations was confirmed by its compatibility with meta-analyses of head circumference (56.49 cm for men and 54.57 cm for women, close to 56.34 cm and 54.49 cm, respectively) and cranial capacity (1282 cm³ for men and 1188 cm³ for women, aligned with 1238 cm³ and 1119 cm³), both compared to the meta-analyses performed.

Although the sample of forensic facial approximations encompasses individuals from multiple continents (Americas, Europe, Africa, and Asia) and distinct chronological periods—extending from the prehistoric era to the 19th century—this distribution remains compatible with the parameters observed in current populations. According to Henneberg (1988), cranial volume in Homo sapiens has not undergone alterations of sufficient magnitude to invalidate comparisons between the Mesolithic and the modern era. In this context, it is observed that contemporary intracultural variability can exceed the variation in means recorded throughout the Holocene [35]. The studies compiled in the present meta-analysis, despite covering different population groups, show a consistent overlap of values and close means, without evidence of relevant outliers when standard deviations are considered (Figures S1–S2). The convergence of data from facial approximations with the other analyzed sets—particularly with the global weighted average—suggests that the values obtained reflect consistent biological tendencies of the species rather than artifacts resulting from sampling variation or geographic distribution.

Although the sample is smaller than the meta-analyses, which included 2348 men and 2074 women for circumference and 3496 men and 3410 women for cranial capacity, it is comparable or superior to peerreviewed studies with smaller populations, such as Allen et al. (2002) (n=23). The validation with 10 actual computed tomography scans, with 7 of the cases between 0 and ±1 SD and 3 cases between +1 and +2 SD (two of which are very close to the 1 SD limit), reinforces the reliability of the triple-measure table (Table 2), which was designed to facilitate clinical consultation with rounded values, maintaining the necessary precision for screening.

Despite its robustness, the study has limitations. The sample of 100 forensic facial approximations, although well distributed, is smaller than meta-analyses, and the unequal proportion (65 men, 35 women) can be seen as a restriction, although it did not compromise the results. The female subsample (n=35) remains within the range commonly reported in morphometric and neuroanatomical studies, including some used in the reference metaanalyses, and therefore does not represent a methodological limitation. Moreover, as the primary objective was to evaluate the relationship between cranial measurements and endocranial volume within a screening framework, internal consistency and compatibility with external parameters are more relevant than strict demographic balance. The observed agreement with meta-analytical values indicates that the dataset adequately represents expected biological variability despite the numerical imbalance. The rounding in the triple table, necessary for clinical simplification, may underestimate borderline cases, such as in the case of microcephaly (-1.93 SD vs. -2.11 SD). However, this simplification does not harm the screening objective, as it signals the need for additional investigation, besides the fact that the online/offline (html) calculator allows for a more precise extraction of data. The pilot test with 10 CT scans, although sufficient for the preliminary objective, includes a case of a large skull that presented a “submerged” brain volume in the region below the ears, not fully reflected in the measurements, but still within the space between +1 SD and +2 SD (Figure 2). This result suggests that the triple measure may not fully capture cases of megalencephaly. This problem was circumvented with the restructuring of the curve based on the pilot cases (Figure S6), which in practice raised the sample from n=100 to n=110; but it can still be better supported if future studies complement the region composed of skulls with high volumetry. Regarding the region composed of neonates, infants, and children, the extrapolation of the curve, based on academic studies that addressed that age group, proved to be very compatible with the pilot cases involving 7 skulls, whose triple measure was performed, reflecting compatibility with projections based on cerebral/cranial volume, age, and head circumference (Figure 3). The pediatric component should be interpreted as an exploratory extension of the model, since the reference values were derived from aggregated literature data rather than individual measurements fully compatible with the proposed method.

Forensic facial approximation demonstrated utility in an atypical approach in this study, combining anthropometry and clinical screening, which suggests that forensic methods can complement medicine, especially in contexts with limited resources.

The potential concern regarding indirect measurements derived from facial approximation and anatomical deformation should be interpreted within the structural constraints of the method. Forensic facial approximation techniques are based on anatomically grounded cranial geometry, and in the present study the measurements were obtained primarily from the cranial vault region, which presents greater morphological stability compared to projection-dependent facial structures. Furthermore, the deformation process is guided by individual cranial dimensions, meaning that the resulting geometry reflects the target morphology rather than donor-specific characteristics. The structural reliability of forensic facial approximation has been previously demonstrated through compatibility between reconstructed faces and identified individuals in forensic contexts [36], while anatomical deformation approaches applied to cranial structures have shown consistency with anatomically derived measurements when compared with tomographic reconstructions in independent studies [37].Beyond their application in forensic facial approximation, the techniques addressed in this study are directly employed in the medical field. Three-dimensional reconstruction based on computed tomography provides models with high geometric fidelity, widely used in surgical planning and structural anatomical analyses. On this basis, digital anatomical deformation is applied as a mechanism of spatial adaptation between structures, guided by morphological correspondence and areas of influence, allowing a reference model to be adjusted to a target anatomy while maintaining structural coherence consistent with the anatomical proportions required for the clinical application. This approach is used in different clinical contexts, including the digital structural prediction of postoperative outcomes in patients with facial deformities [38], while photogrammetry is applied for similar predictive purposes in rhinoplasty planning [39]. The integration of photogrammetry with anatomical deformation is also employed in the fabrication of facial prostheses for oncological patients, a process that requires submillimetric precision [40].

Together, these factors support the validity of the present methodological framework for volumetric estimation purposes.

For future research, validation of the table with larger and more diverse samples, including diagnosed cases of microcephaly and megalencephaly, is recommended to confirm its sensitivity across the entire spectrum of cranial–brain deviations. Integration with magnetic resonance imaging can increase accuracy in advanced clinical settings.

One point to be evaluated is that the present work made measurements from digital models, without the volume of the hair, which, while not compressing the soft tissue as in real measurements, could lead to an underestimation of the measurements, given the potential increase from the hair group. Future studies could evaluate, in real cases versus CT scans or even lateral radiographs, what the margin of error should be established for greater statistical confidence.

It is important to emphasize that the application of the triple measure is non-invasive, poses no risks to the individual, and requires minimal effort from the specialist, as the measurements are simple and quick. Furthermore, it works even with the absence of one of the three data points, meaning the survey remains valid even if problems occur during the process. Another important factor is the high accessibility of the solution, since, thanks to the printed table, it is possible to access the data without the need for a digital tool and, even in this case, there is no need for an internet connection, since the calculator is an html/javascript page that runs on any PC or smartphone operating system, simply by downloading the page and running it, practically, as a native app.

The social impact of the table is significant, especially in vulnerable populations, such as those affected by the Zika virus. Early screening can improve the quality of life, reducing complications associated with microcephaly, such as intellectual disability and epilepsy. The tool, therefore, has the potential to contribute to public health and medicine, but its implementation requires additional testing to ensure effectiveness and reliability in different clinical and population contexts.

## Conclusion

The present study introduced and evaluated an anthropometric model based on the triple measure — composed of head circumference, sagittal arc, and coronal arc — as a complementary approach for estimating endocranial volume. The results demonstrated that the triple measure shows a more consistent association with endocranial volume than head circumference alone, while maintaining operational simplicity and a non-invasive nature. Additional validation analyses indicated good agreement between estimated and measured values, with reduced prediction errors compared to the traditional method based solely on head circumference.

The proposed model is not intended to replace diagnostic methods based on neuroimaging, but rather to serve as an accessible anthropometric screening tool capable of assisting in the identification of potential deviations from expected patterns and, when necessary, indicating the need for complementary examinations. In this context, its applicability may be particularly relevant in settings with limited access to imaging resources.

Despite the promising findings, the study presents limitations related to sample size, heterogeneity of data sources, and the exploratory nature of the pediatric extension, which was constructed from aggregated literature data. Future investigations with larger samples, including individuals with confirmed diagnoses and data obtained directly from neuroimaging, may improve clinical robustness and validate the model’s performance across different populations.

Overall, the findings suggest that the triple measure constitutes a plausible and potentially useful approach for the initial screening of cranial–brain patterns, providing a foundation for future research and possible complementary clinical applications.

## Supplementary Material

Additional resources supporting the findings of this study, including extended data tables, detailed validation parameters, and the digital calculation tool, are available for review at the following URL: https://docs.google.com/document/d/15H3QQbHE_ptJ01gvOWbjekR7LMPPxn3x/edit?usp=sharing

## Ethical approval and consent to participate

This study used anonymized retrospective imaging data and publicly available digital models. No identifiable personal data were accessed. According to institutional and national regulations, formal ethics committee approval and informed consent were not required.

## Author contributions

C.M.: Conceptualization, methodology, software development, data curation, formal analysis, visualization, writing – original draft.

J.Y.A.: Methodological supervision, clinical interpretation, writing – review & editing.

A.B.P.: Historical and anatomical contextualization, writing – review & editing.

A.S.: Anatomical analysis, provision of comparative cranial material, writing – review & editing.

J.Š.: Technical support, provision of cranial data and materials, writing – review & editing.

I.K.: Anatomical expertise, provision of cranial material, writing – review & editing.

E.V.: Archaeological context and interpretation, provision of skeletal material, writing – review & editing.

T.B.: Dental and craniofacial interpretation, writing – review & editing.

F.M.G.: Conceptual discussion, anatomical validation, supervision, writing – review & editing.

All authors reviewed and approved the final manuscript.

## Funding

This research did not receive any specific grant from funding agencies in the public, commercial, or not-forprofit sectors.

## Notes

https://docs.google.com/document/d/15H3QQbHE_ptJ01gvOWbjekR7LMPPxn3x/edit?usp=sharing

### Competing Interest Statement

The authors have declared no competing interest.

## References

[1] Souza WC. A importância da antropometria no ambiente escolar. REBESDE. 2022;3(2).

[2] Madureira NL. A estatística do corpo: antropologia física e antropometria na alvorada do século XX. Etnográfica. 2003;7(2):283–303. 10.4000/etnografica.2899

[3] Macchiaverni LML, Barros Filho AA. Perímetro cefálico: por que medir sempre. Medicina (Ribeirão Preto). 1998;31(4):595–609.

[4] Giedd JN. The teen brain: insights from neuroimaging. J Adolesc Health. 2008;42(4):335–343. 10.1016/j.jadohealth.2008.01.007

[5] Moraes C, Habicht ME, Galassi FM, Varotto E, Beaini T. Pharaoh Tutankhamun: a novel 3D digital facial approximation. Ital J Anat Embryol. 2023;127(1):13–22. 10.36253/ijae-14514

[6] Hanzlik E, Gigante J. Microcephaly. Children. 2017;4(6):47. 10.3390/children4060047

[7] Hofman MA. A biometric analysis of brain size in micrencephalics. J Neurol. 1984;231(2):87–93. 10.1007/BF00313723

[8] Peixoto HM, Gallo LG, Nascimento GL, Oliveira MRF. Economic impact of Zika virus infection and associated conditions. In: Zika virus biology, transmission, and pathology. Elsevier; 2021. p. 63–74. 10.1016/B978-0-12-820268-5.00006-0

[9] Lee J. Malformations of cortical development: genetic mechanisms and diagnostic approach. Korean J Pediatr. 2017;60(1):1–9. 10.3345/kjp.2017.60.1.1

[10] Pirozzi F, Nelson B, Mirzaa G. From microcephaly to megalencephaly: determinants of brain size. Dialogues Clin Neurosci. 2018;20(4):267–282. 10.31887/dcns.2018.20.4/gmirzaa

[11] Moraes CAC, Dias PEM, Melani RFH. Demonstration of protocol for computer-aided forensic facial reconstruction with free software and photogrammetry. J Res Dent. 2014;2(1):77–90. 10.19177/jrd.v2e1201477-90

[12] Abdullah JY, Moraes C, Saidin M, Rajion ZA, Hadi H, Shahidan S, Abdullah JM. Forensic facial approximation of a 5000-year-old female skull from a shell midden in Guar Kepah, Malaysia. Appl Sci. 2022;12(15):7871. 10.3390/app12157871

[13] Quatrehomme G, Cotin S, Subsol G, Delingette H, Garidel Y, Grévin G, et al. A fully three-dimensional method for facial reconstruction based on deformable models. J Forensic Sci. 1997;42(4):649–652.

[14] Lee JJ, McGue M, Iacono WG, Michael AM, Chabris CF. The causal influence of brain size on human intelligence: evidence from within-family phenotypic associations and GWAS modeling. Intelligence. 2019;75:48–58. 10.1016/j.intell.2019.01.011

[15] Bale SJ, Amos CI, Parry DM, Bale AE. Relationship between head circumference and height in normal adults and in the nevoid basal cell carcinoma syndrome and neurofibromatosis type I. Am J Med Genet. 1991;40(2):206–210. 10.1002/ajmg.1320400217

[16] Bushby KM, Cole T, Matthews JN, Goodship JA. Centiles for adult head circumference. Arch Dis Child. 1992;67(10):1286–1287. 10.1136/adc.67.10.1286

[17] Catapan MF, Okimoto ML, Villas Boas M, Waldhauer R. Statistical analysis of the height of the human head in the use of ballistic helmets. In: Ahram T, Karwowski W, Marek T, editors. Proc 5th Int Conf Appl Hum Factors Ergon (AHFE 2014). AHFE; 2014. p. 1–11.

[18] Costa NR, Mancine L, Salvini R, Teixeira JM, Rodriguez RD, Leite REP, et al. Microcephaly measurement in adults and its association with clinical variables. Rev Saude Publica. 2022;56:38. 10.11606/s1518-8787.2022056004175

[19] Haeri SMJ, Vakilian K, Moslemi A, Maleki M, Babazadeh-Zavieh SS, Bayat M. Relationship between parental head circumference and birth type: a pilot study. Open Public Health J. 2023;16(1). 10.2174/0118749445268235231117080517

[20] Khursheed Z, Itoo MS, Iqbal Z, Bhat GM. Relationship between height and head circumference in Kashmiri population. J Cardiovasc Dis Res. 2024;15(7):2064–2071.

[21] Mansur DI, Haque MK, Sharma K, Mehta DK, Shakya R. Use of head circumference as a predictor of height of individual. Kathmandu Univ Med J. 2014;12(46):89–92.

[22] Örmeci AR, Gürbüz H, Ayata A, Çetin H. Adult head circumferences and centiles. J Turgut Özal Med Cent. 1997;4(3):261–264.

[23] Shahabuddin, Haq AU, Khushbakht U, Siddiqui N. Estimation of height using head circumference of medical students in Peshawar, Pakistan. J Postgrad Med Inst. 2024;38(1):64–69. 10.54079/jpmi.38.1.3289

[24] Allen JS, Damasio H, Grabowski TJ. Normal neuroanatomical variation in the human brain: an MRI-volumetric study. Am J Phys Anthropol. 2002;118(4):341–358. 10.1002/ajpa.10092

[25] Courchesne E, Chisum HJ, Townsend J, Cowles A, Covington J, Egaas B, et al. Normal brain development and aging: quantitative analysis at in vivo MR imaging in healthy volunteers. Radiology. 2000;216(3):672–682. 10.1148/radiology.216.3.r00au37672

[26] Panta OB, Gurung B, Giri SR, Adhikari A, Ghimire RK. Mean intracranial volume of brain among patients with normal magnetic resonance imaging referred to a tertiary care centre. J Nepal Med Assoc. 2023;61(268):934–937. 10.31729/jnma.8357

[27] Peters M, Jäncke L, Staiger JF, Schlaug G, Huang Y, Steinmetz H. Unsolved problems in comparing brain sizes in Homo sapiens. Brain Cogn. 1998;37(2):254–285. 10.1006/brcg.1998.0983

[28] Ritchie SJ, Cox SR, Shen X, Lombardo MV, Reus LM, Alloza C, et al. Sex differences in the adult human brain: evidence from 5,216 UK Biobank participants. Cereb Cortex. 2018;28(8):2959–2975. 10.1093/cercor/bhy109

[29] Shepur MP, Magi M, Nanjundappa B, Havaldar PP, Gogi P, Saheb SH. Morphometric analysis of endocranial capacity. Int J Anat Res. 2014;2(1):242–248.

[30] Bartholomeusz HH, Courchesne E, Karns CM. Relationship between head circumference and brain volume in healthy normal toddlers, children, and adults. Neuropediatrics. 2002;33(5):239–241. 10.1055/s-2002-36735

[31] Burkhardt W, et al. Non-invasive estimation of brain volume in infants. Early Hum Dev. 2019;132:52–57. 10.1016/j.earlhumdev.2019.03.020

[32] Gerber AJ, et al. Anatomical brain magnetic resonance imaging of typically developing children and adolescents. J Am Acad Child Adolesc Psychiatry. 2009;48(5):465–470. 10.1097/CHI.0b013e31819f2715

[33] Knickmeyer RC, Gouttard S, Kang C, Evans D, Wilber K, Smith JK, et al. A structural MRI study of human brain development from birth to 2 years. J Neurosci. 2008;28(47):12176–12182. 10.1523/JNEUROSCI.3479-08.2008

[34] McAllister A, et al. Quantitative synthetic MRI in children: normative intracranial tissue segmentation values during development. AJNR Am J Neuroradiol. 2017;38(12):2364–2372. 10.3174/ajnr.A5398

[35] Henneberg, M. (1988). Decrease of Human Skull Size in the Holocene. Human Biology, 60(3), 395–405. http://www.jstor.org/stable/41464021

[36] Baldasso, R. P., Moraes, C., Gallardo, E., Stumvoll, M. B., Crespo, K. C., Strapasson, R. A. P., & de Oliveira, R. N. (2020). 3D forensic facial approximation: Implementation protocol in a forensic activity. Journal of Forensic Sciences, 66(1), 383–388. 10.1111/1556-4029.14587

[37] Moraes, C. (2024). A Aproximação Facial Forense de Jebel Irhoud (∼315.000 AP). figshare. 10.6084/M9.FIGSHARE.26038636

[38] Lobo, F., Filho, L. I., Sigua-Rodriguez, E. A., da Silva, B. G., Tolentino, E. de S., Borges, Y. M., da Silva, M. C., Tonin, R. H., & Iwaki, L. C. V. (2022). Evaluation of ortogonblender software bone movement tools in bimaxillary orthognatic surgeries performed in dolphin software. In Journal of Stomatology, Oral and Maxillofacial Surgery (Vol. 123, Issue 4, pp. 417–421). Elsevier BV. 10.1016/j.jormas.2021.10.001

[39] Sobral, D. S., Duarte, D. W., Dornelles, R. F. V., & Moraes, C. A. C. (2021). 3D Virtual Planning for Rhinoplasty Using a Free Add-On for Open-Source Software. Aesthetic Surgery Journal, 41(8), NP1024–NP1032. 10.1093/asj/sjab085

[40] Salazar-Gamarra, R., Cárdenas-Bocanegra, A., Masch, U., Da Costa Moraes, C. A., Seelaus, R., Lopes Da Silva, J. V., & Lauria Dib, L. (2022). Color translation from monoscopic photogrammetry +ID Methodology into a Polyjet final 3D printed facial prosthesis. F1000Research, 11, 582. 10.12688/f1000research.111196.1

